# Multiple Nested Distributed Language Networks in the Human Brain

**DOI:** 10.64898/2026.08.07.743553

**Authors:** Jingnan Du, Anne Billot, Wendy Sun, Gregory Hickok, Mark Eldaief, Randy L. Buckner

**Author notes:** Correspondence (R.L.B.).

## Abstract

Brain regions specialized for language have been extensively described, yet their arrangement into one or multiple networks remains debated. Using precision functional mapping across three independent cohorts of intensively scanned individuals (22 individuals scanned over 216 separate MRI sessions), we dissociated two nested left-lateralized perisylvian networks: an intermediate language network (intLANG) and an anatomically distinct association language network (aLANG). intLANG is anchored to precentral speech areas and the Sylvian parietal-temporal area (Spt), whereas aLANG surrounds intLANG and extends into higher-order prefrontal and temporal association cortices. The two networks can be fully recapitulated by functional connectivity from adjacent cerebellar regions, indicating that they are segregated, brain-wide networks. Task-based analyses further reveal that intLANG and aLANG are functionally distinct: intLANG responds robustly during rhyme judgments and nonword reading that emphasize phonology, whereas aLANG is preferentially recruited during meaning-based sentence processing. These findings indicate that human language engages nested distributed networks each specialized for distinct components of language processing: a lower-order network biased toward phonology, and a surrounding association network that subserves higher-order syntax and semantics. This nested organization is similar to other brain systems suggesting a shared hierarchical motif that may give rise to specialized cognitive functions across the human brain.

## Background

Human language is supported by distributed left-lateralized regions situated around the Sylvian fissure that include, but are not limited to, classical regions known as Broca’s and Wernicke’s areas (Geschwind 1965; Friederici 2011; Hickok 2025). While there is general agreement on the existence of multiple specialized regions within these broad cortical zones, there is ongoing debate about how they are organized into networks and the nature of functional specializations that differentiate them. A fundamental debate centers on whether there is a single distributed network that serves as the core integrator for linguistic functions or whether there are multiple interacting networks, each specialized for distinct components of language.

One influential framing proposes that a large, distributed network – referred to as the language network – supports multiple linguistic computations by interacting with speech perception and articulatory motor zones (Fedorenko et al. 2024). Consistent with historical expectations, the topography of the monolithic language network includes regions at and around classically defined Broca’s and Wernicke’s areas. This network was identified by mapping brain regions that increase their response when participants read or hear meaningful sentences (Fedorenko et al. 2010; see also Friederici et al. 2000; Vandenberghe et al. 2002; Humphries et al. 2006; Price 2012); it is robustly activated by sentence processing tasks across a range of natural and constructed languages (Malik-Moraleda et al. 2022,2025); and the distributed pattern of the language network can be recapitulated by intrinsic functional connectivity, suggesting that it may be supported by a stable anatomically connected network (Braga et al. 2020; Du et al. 2024; Salvo et al. 2025; Anderson et al. 2026).

A prominent alternative theory is that the neural architecture of language is organized as a nested hierarchy where the levels of the hierarchy correspond to the major linguistic computational systems: roughly, sound structure (phonology) organized at a lower level closer to sensory and motor systems and sentence structure (syntax) organized at a higher level closer to, and interacting with, the semantic system (Hagoort 2013; Friederici 2017; Hickok 2025). By this view, language is not supported by a single monolithic network, but rather multiple networks that are differentially specialized for distinct levels of linguistic processing.

A challenge in differentiating these two models is that both converge on the same general group of perisylvian language zones – it is primarily the delineation and arrangement of closely juxtaposed regions into networks that is postulated to differ. One critical difference is that in the monolithic model, all interactions between anterior and posterior language zones are through the core interconnected association network (Fedorenko et al. 2024). By contrast, the hierarchical model proposes that distinct parietal-temporal regions interact with distinct frontal regions through partially segregated pathways; each pathway preferentially processes a separate type of linguistic information. In a prior set of studies, we noted a small, distributed network that is fragmented and variable across individuals, yet intriguingly positioned between the early motor / auditory regions and the canonical association language network (Braga et al. 2020).

Here, using precision within-subject functional neuroimaging, we provide strong evidence that language function is supported by multiple nested distributed networks. We identified a lower-order network, labeled the intermediate language network (intLANG), involving posterior regions near auditory cortex, including Spt, and frontal regions near the laryngeal and orofacial motor representations (Bouchard et al. 2013; Eichert et al. 2020). A distinct higher-order network, labeled the association language network (aLANG) surrounds intLANG with regions extending into parietal-temporal and prefrontal association cortex. The two networks – intLANG and aLANG – are anatomically adjacent with juxtaposed component regions in both anterior and posterior cortex, making their separation challenging. On inspection, the intLANG resembles aspects of the dorsal stream phonological system anticipated by the dual stream model (Hickok and Poeppel 2007; Hickok 2025), while the aLANG converges with the network emphasized by Fedorenko and colleagues (2010, 2024) and the syntactic network identified by others (see Matchin and Hickok, 2020, for a review). Once the two networks were identified as anatomically distinct – a result replicated across three independent studies – their functional response properties could be robustly dissociated along linguistic dimensions.

## Results

### Overview

We characterized the organization of human language networks across three independent, intensively sampled participant cohorts. Study 1 participants were initially examined to identify intLANG using functional connectivity within the idiosyncratic anatomy of each individual. intLANG reliably included multiple frontal regions at or near the precentral gyrus (labeled the dorsal and ventral precentral speech areas, dPCSA and vPCSA, respectively; Hickok et al. 2023) and the anatomically distant posterior parietal-temporal region, Spt. A second network, aLANG, was identified surrounding intLANG and extending into association cortices. Both intLANG and aLANG were left-lateralized. To establish that the two networks are partially segregated brain-wide networks, functional connectivity was investigated from the contralateral cerebellum. Both intLANG and aLANG, including their distributed anterior and posterior components, could be fully recapitulated by placing seed regions in closely adjacent subdivisions of the cerebellum.

Having established that the two networks are anatomically separate, we next examined their functional response properties. intLANG was preferentially recruited by phonological processing in two separate contrasts: Rhyme > Face from an N-Back working memory task and Nonword > Fixation from a Sentence Processing task, whereas aLANG showed minimal response. The anatomical and functional dissociations between the two networks were replicated in Studies 2 and 3. As a final analysis to confirm a functional double dissociation, the Nonword > Fixation contrast, which preferentially demands phonological processing, was directly compared to the Sentence > Nonword contrast, which has been a widely used localizer that engages higher-order syntactic and semantic processing. A robust functional double dissociation between the two networks emerged, establishing that intLANG and aLANG are anatomically separate, nested networks supporting distinct linguistic processes.

### intLANG can be identified and replicated within the idiosyncratic anatomy of individuals

Motivated by the possibility that an intermediate network might exist near the precentral speech regions (Bouchard et al. 2013; Eichert et al. 2020; Silva et al. 2022), we began our investigations by first localizing the tongue / mouth motor representation within each individual using a motor task or, as backup, functional connectivity (Saadon-Grosman et al. 2022; Du et al. 2024). Then seed regions were manually placed along the precentral gyrus anterior to the tongue / mouth representation to determine whether a distributed network could be revealed. Based on prior reports, we anticipated that such a network would include dorsal and ventral precentral components and a posterior component near the parietal-temporal junction. We focused initially on the five participants in Study 1 who each had enough data to be separated into independent within-individual datasets, with the remaining participants from Studies 2 and 3 set aside for prospective replication. In each Study 1 participant, the seed regions revealed a distributed left-lateralized network that sat anatomically between the tongue / mouth motor representation and the extended association language network, echoing observations our laboratory has made previously (Braga et al. 2020; see also Glasser et al. 2016). In almost all individuals, the network comprised the anticipated dorsal and ventral precentral regions, and a posterior region falling at or near the parietal-temporal junction, together with a frontal midline region likely localized to the supplementary motor area (SMA).

To establish that these separate regions belong to a unified network, we next systematically placed seed regions in each of the separate distributed zones (paralleling Braga et al. 2020). The same general pattern was recovered from each region. That is, the precentral regions could be localized by placing seed regions in Spt or SMA, and the posterior Spt region could be localized by placing seed regions in SMA, dPCSA or vPCSA. Given that seed regions across all of the distributed locations recover the same network, the midline SMA region was chosen to display the network because it allows the topography of the network regions along the precentral gyrus and posterior cortical language zones to be visualized in a spatially unbiased manner, without the local correlation that surrounds the seed region. intLANG reliably included regions at or near dPCSA, vPCSA, and Spt (Fig. 1; see group-level estimates in Hickok et al. 2023).

**Figure 1.**
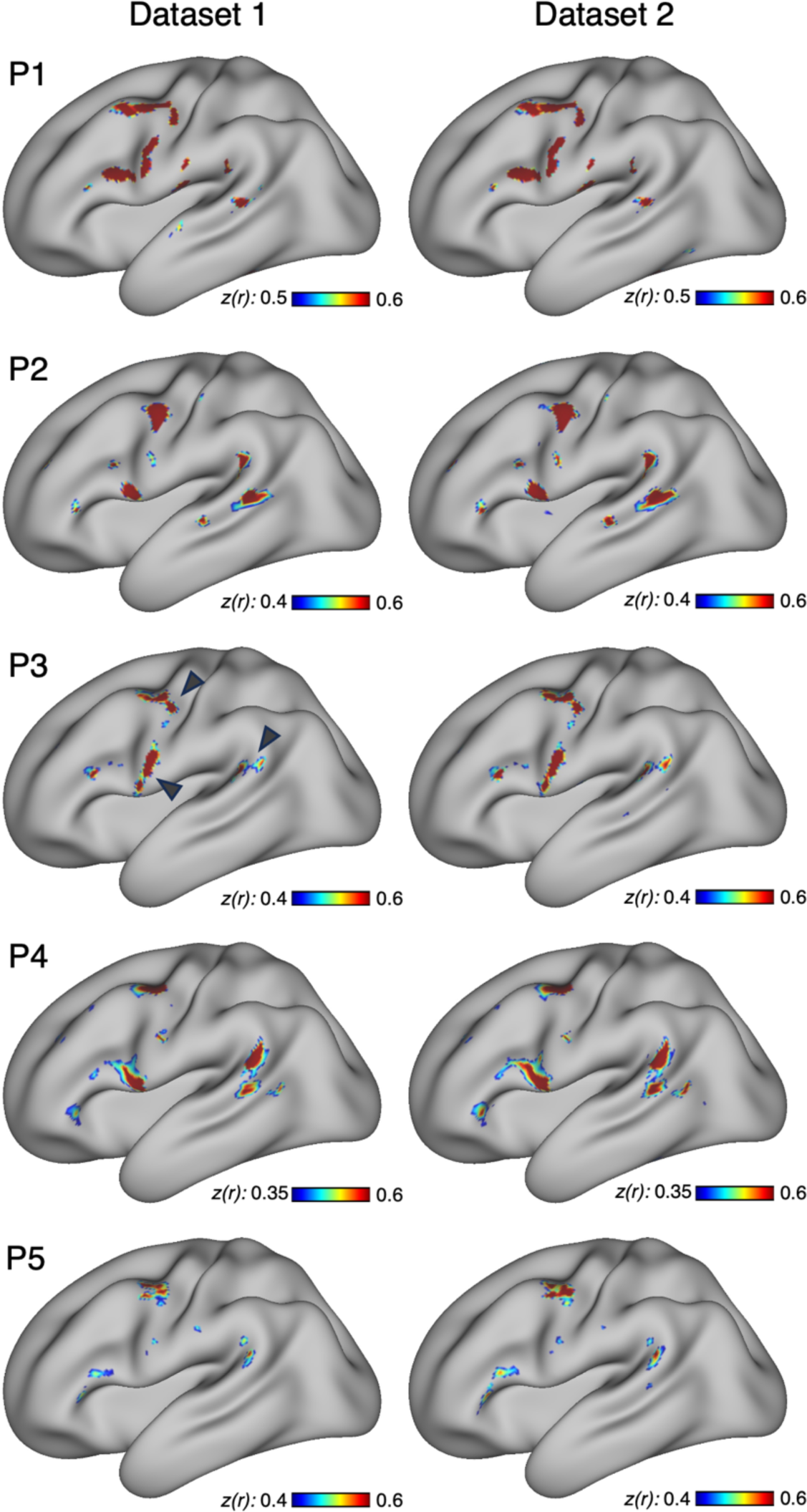
The intermediate language network (intLANG) can be identified and replicated within individuals. Intrinsic functional connectivity maps are displayed on the inflated left-hemisphere cortical surface for five participants from Study 1 (P1–P5). For each participant, two independent datasets are shown in the two columns (Dataset 1 and Dataset 2) which are replications of the connectivity patterns within the idiosyncratic anatomy of each participant. The displayed intLANG estimate for each participant used a single-vertex seed region placed at or near the supplementary motor area (SMA) to provide for a spatially unbiased visualization of the regions on the lateral surface. Similar maps are produced from seed regions placed within any of the component regions (see Supplementary Figures). In each individual, intLANG included regions at or near the dorsal precentral speech area (dPCSA), the ventral precentral speech area (vPCSA), and the Sylvian parietal-temporal area (Spt). Black arrowheads point to the locations of these three anchor regions in P3. Correlation maps are plotted as *z(r)* using the JET256 color scale.

Bilateral surfaces are shown in the Supplemental Materials. intLANG was left-lateralized. On average it occupied 2.01% of left-hemisphere cortical vertices but only 0.12% of right-hemisphere vertices. The lateralization index averaged 0.91 (0.74 - 1.00), with a positive value in every participant.

Spatial variability existed between participants. While the general network pattern was consistent, the detailed topography varied idiosyncratically (Supplemental Materials). dPCSA fell along the precentral gyrus ventral to the superior frontal sulcus, but sometimes extending anteriorly. vPCSA was localized just dorsal to the Sylvian fissure. The precise extent of the Spt region differed across individuals but generally fell in the posterior portion of the Sylvian fissure at the parietal-temporal junction. Several participants possessed a second posterior region ventral to Spt in the superior temporal sulcus (STS), consistent with reports on the initial identification of area Spt (Buchsbaum et al. 2001; Hickok et al. 2023).

Finally, to establish that these idiosyncratic spatial details are reliable, the full spatial pattern of the network was replicated in the independent held-out dataset within each individual (Fig. 1, Dataset 2). The two replicate estimates converged on largely the same set of regions in each individual, despite the spatial differences between individuals, suggesting the idiosyncratic features are stable aspects of within-subject functional anatomy.

### intLANG is spatially juxtaposed with distinct network**s**

To contextualize the anatomy of the intLANG network, we next examined how intLANG was situated relative to previously described networks and regions that are spatially adjacent: the canonical association language network, aLANG (Fedorenko et al. 2010, 2024), given its role in language; the action-mode network (AMN), given its position in and around the precentral gyrus (Gordon et al. 2023; Dosenbach et al. 2025); and the tongue motor representation (Tongue) that provides an anchor to understand intLANG’s position relative to a mapped motor effector region (Saadon-Grosman et al. 2022). Across all five participants, intLANG occupied cortical regions that sat adjacent to, or at the edge of, each of these networks and regions (Fig. 2).

**Figure 2.**
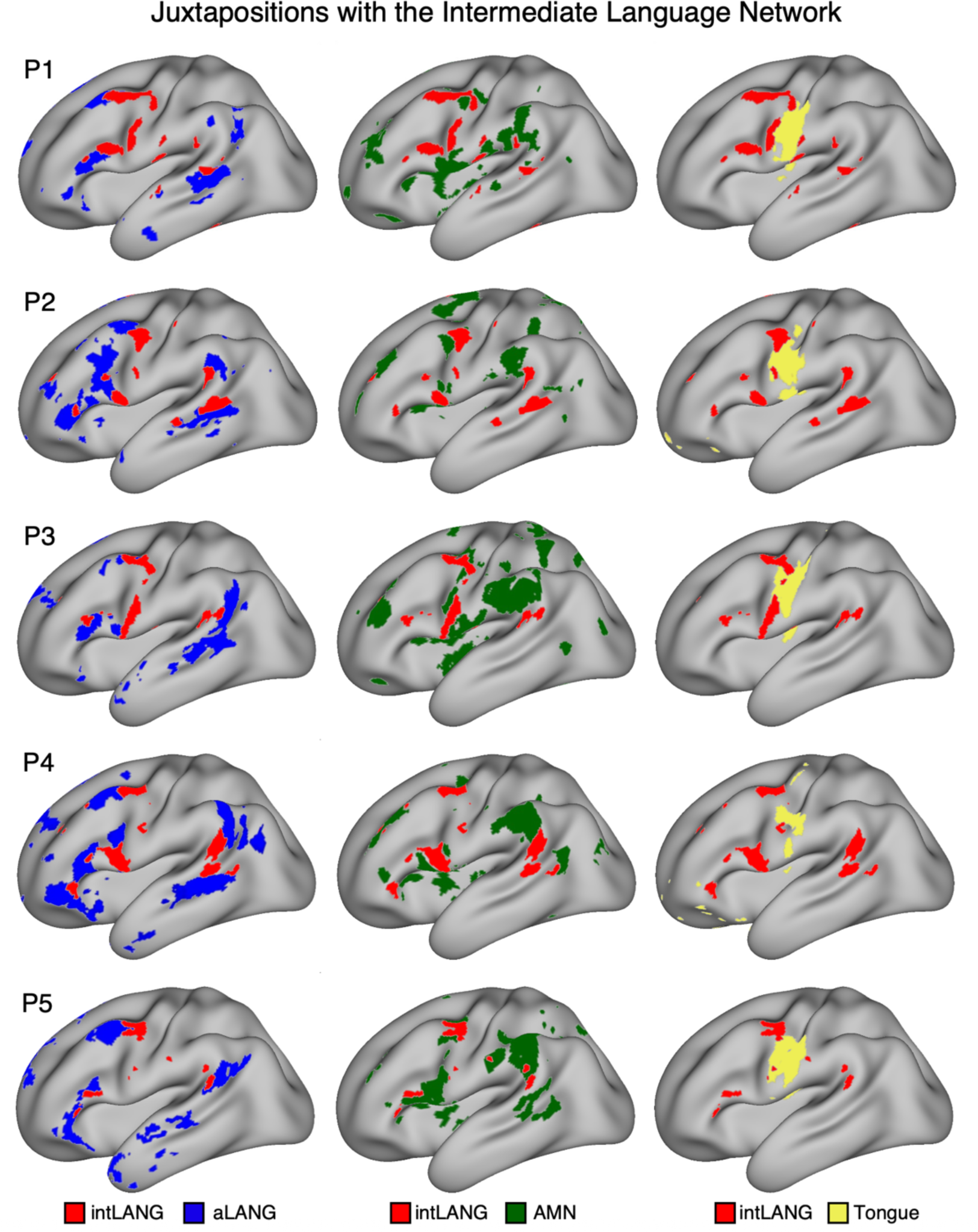
Spatial relationships between the intermediate language network (intLANG) and adjacent networks. Inflated left-hemisphere cortical surfaces are displayed for the five Study 1 participants (P1–P5). intLANG (red) serves as the spatial anchor across all three columns. (**Left column**) intLANG is shown alongside the association language network (aLANG, blue). The two networks exhibited extensive adjacency along the inferior frontal gyrus, the precentral gyrus, and middle-to-posterior superior temporal sulcus, but occupied distinct cortical territories in all participants. (**Middle column**) intLANG is shown alongside the action-mode network (AMN, green). AMN regions were localized between or directly adjacent to intLANG regions, particularly within the inferior and middle frontal gyri, as well as near to the superior temporal sulcus and inferior parietal lobule. **(Right column**) intLANG is shown alongside the Tongue region (yellow), defined from a tongue-movement task contrast or seed-region-based functional connectivity. intLANG regions at or near the dorsal and ventral precentral speech areas (dPCSA and vPCSA) were located adjacent to the Tongue region.

As expected, given how the network was identified, intLANG was anterior to the coarsely mapped Tongue region (Fig. 2, right column). In each individual, intLANG included both dorsal and ventral regions, forming separate dPCSA and vPCSA representations surrounding the Tongue region, consistent with the topographical organization of the precentral gyrus established by intracranial mapping (Bouchard et al. 2013; see also Eichert et al. 2020). In some individuals the ventral region, vPCSA, was minimal (P5) or included a small, interspersed region (P3).

The relationship between intLANG and aLANG (Fig. 2, left column) was characterized by extensive adjacency along the inferior frontal gyrus (IFG), the precentral gyrus, and middle-to-posterior STS. The precise pattern varied across participants, though close spatial juxtapositions were present in every case (see P2 and P4 in Fig. 2 for examples). A similar pattern was observed between intLANG and AMN (Fig. 2, middle column). AMN regions were positioned between or adjacent to intLANG regions, particularly along the IFG and the STS extending into the inferior parietal lobule.

These collective findings show that intLANG is separate from aLANG with a nested spatial relationship: intLANG falls closest to the central sulcus and early auditory cortex, with aLANG surrounding intLANG and extending into prefrontal and temporal association cortices.

### intLANG and aLANG are anatomically distinct networks

A key prediction of the hierarchical model, distinguishing it from the monolithic model of language network organization, is the existence of distinguishable distributed networks. The prediction is that regions associated with intLANG will be more correlated with each other, including spatially distant regions, than with aLANG regions, including those that are spatially adjacent. To test this prediction, we identified component regions of both intLANG and aLANG in each participant’s Dataset 1 and then estimated the region-to-region correlation matrix in each participant’s Dataset 2, ensuring an unbiased test of whether intLANG and aLANG segregate.

The functional connectivity matrices revealed a block-modular organization consistent with segregated distributed networks (Fig. 3). The three intLANG regions (dPCSA, vPCSA, and Spt) were strongly correlated with one another, forming a block in the upper-left of the matrix. The five aLANG regions (IFG, posterior middle frontal gyrus [pMFG], anterior superior temporal sulcus [aSTS], middle superior temporal sulcus [mSTS], and posterior superior temporal sulcus [pSTS]) were strongly correlated with one another, forming a block in the lower-right. By contrast, correlations between the intLANG and aLANG regions, occupying the off-diagonal blocks, were consistently weaker.

**Figure 3.**
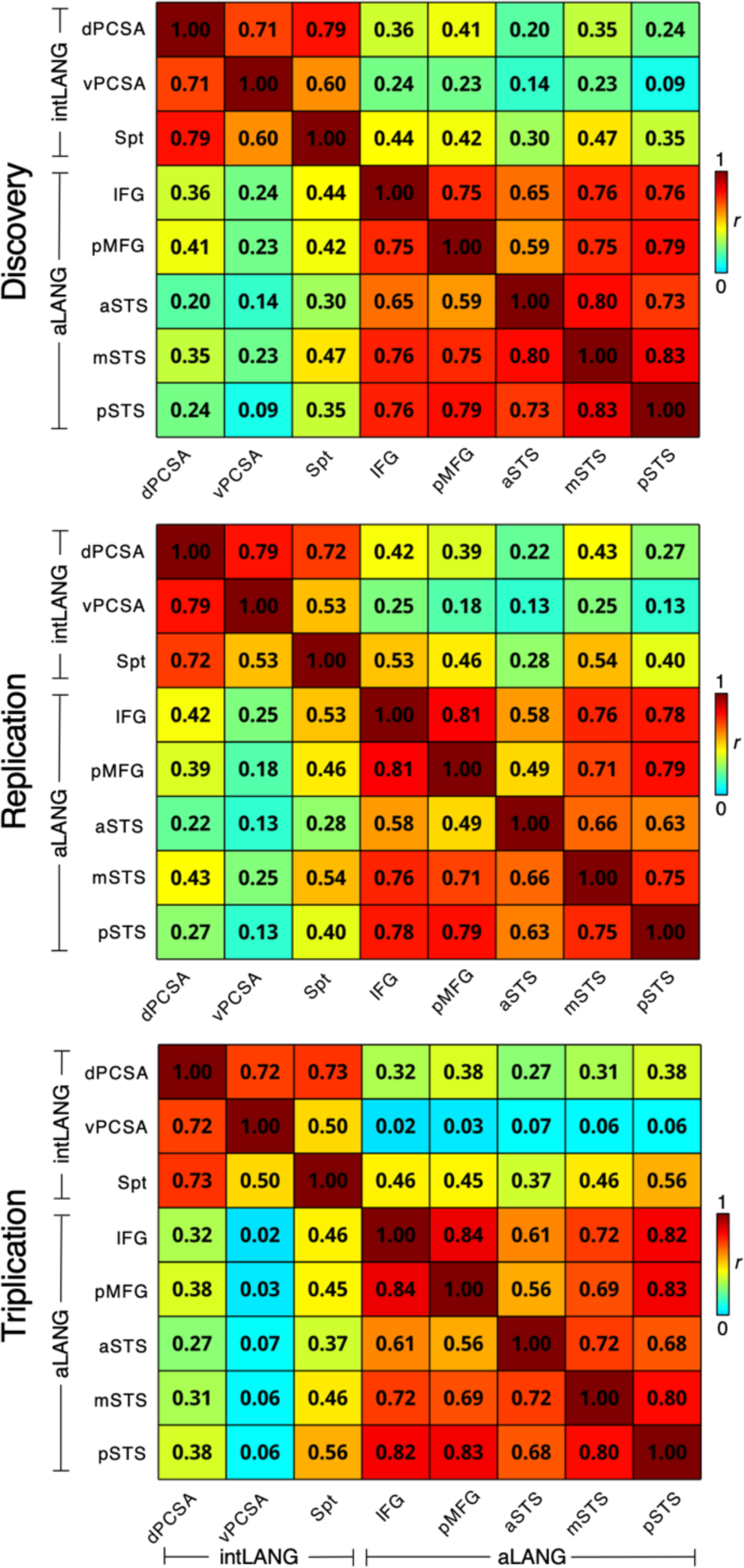
Segregation of the intermediate language network (intLANG) and the association language network (aLANG) is reproducible. Functional connectivity matrices show the pairwise Pearson correlations across three intLANG and five aLANG regions. The matrices consistently revealed a block-modular organization: the three intLANG regions (upper-left 3 × 3 block) were strongly correlated with one another, the five aLANG regions (lower-right 5 × 5 block) were strongly correlated with one another, and the between-network correlations were weaker (off-diagonal blocks). intLANG regions included the dorsal precentral speech area (dPCSA), ventral precentral speech area (vPCSA), and Sylvian parietal-temporal area (Spt); aLANG regions included the inferior frontal gyrus (IFG), posterior middle frontal gyrus (pMFG), anterior superior temporal sulcus (aSTS), middle superior temporal sulcus (mSTS), and posterior superior temporal sulcus (pSTS). Matrices are displayed for each cohort separately (Studies 1-3, designated Discovery, Replication, and Triplication). Within each cohort, regions were defined within each individual using Dataset 1, and pairwise correlations were computed from the independent Dataset 2. Functional connectivity matrices were Fisher *z*-transformed, averaged across participants within each cohort, and converted back to *r* values for visualization. The color scale indicates the strength of functional connectivity.

Post-hoc statistical analyses confirmed the separation between networks. The mean within-intLANG functional connectivity was r = 0.71 (*z(r)* = 0.88) and the mean within-aLANG functional connectivity was r = 0.75 (*z(r)* = 0.97), each substantially greater than the mean between-network functional connectivity of r = 0.30 (*z(r)* = 0.31). Both differences were significant even considering the small sample size (within-intLANG > between, *t*(4) = 6.40, *p* < 0.01, Cohen’s *d* = 2.86; within-aLANG > between, *t*(4) = 9.47, *p* < 0.001, Cohen’s *d* = 4.24). These patterns were replicated in the larger, independent cohorts in Studies 2 and 3.

### intLANG and aLANG are segregated brain-wide networks

If intLANG and aLANG are segregated brain-wide networks, their separation should be echoed in structures beyond the cerebral cortex, including the contralaterally organized cerebellum (e.g., Xue et al. 2021; Saadon-Grosman et al. 2024; Casto et al. 2026). Given this possibility, we explored whether intLANG and aLANG possess spatially separate cerebellar representations and further whether seed regions placed within the cerebellum can recover the distinct cerebral cortical networks.

Fig. 4 shows the results for one participant (P2) with the remaining Study 1 participants presented in Supplemental Materials. The full extent of the cerebral intLANG network could be recapitulated and distinguished from the aLANG network by placing adjacent seed regions at or near Crus I/II and Lobule V and, separately, by placing adjacent seed regions at or near Lobule VIII. This double representation is predicted by, and consistent with, the known functional organization of the cerebellum, where all mapped networks have distinct anterior and posterior representations (e.g., Buckner et al. 2011; Guell et al. 2018; Xue et al. 2021).

**Figure 4.**
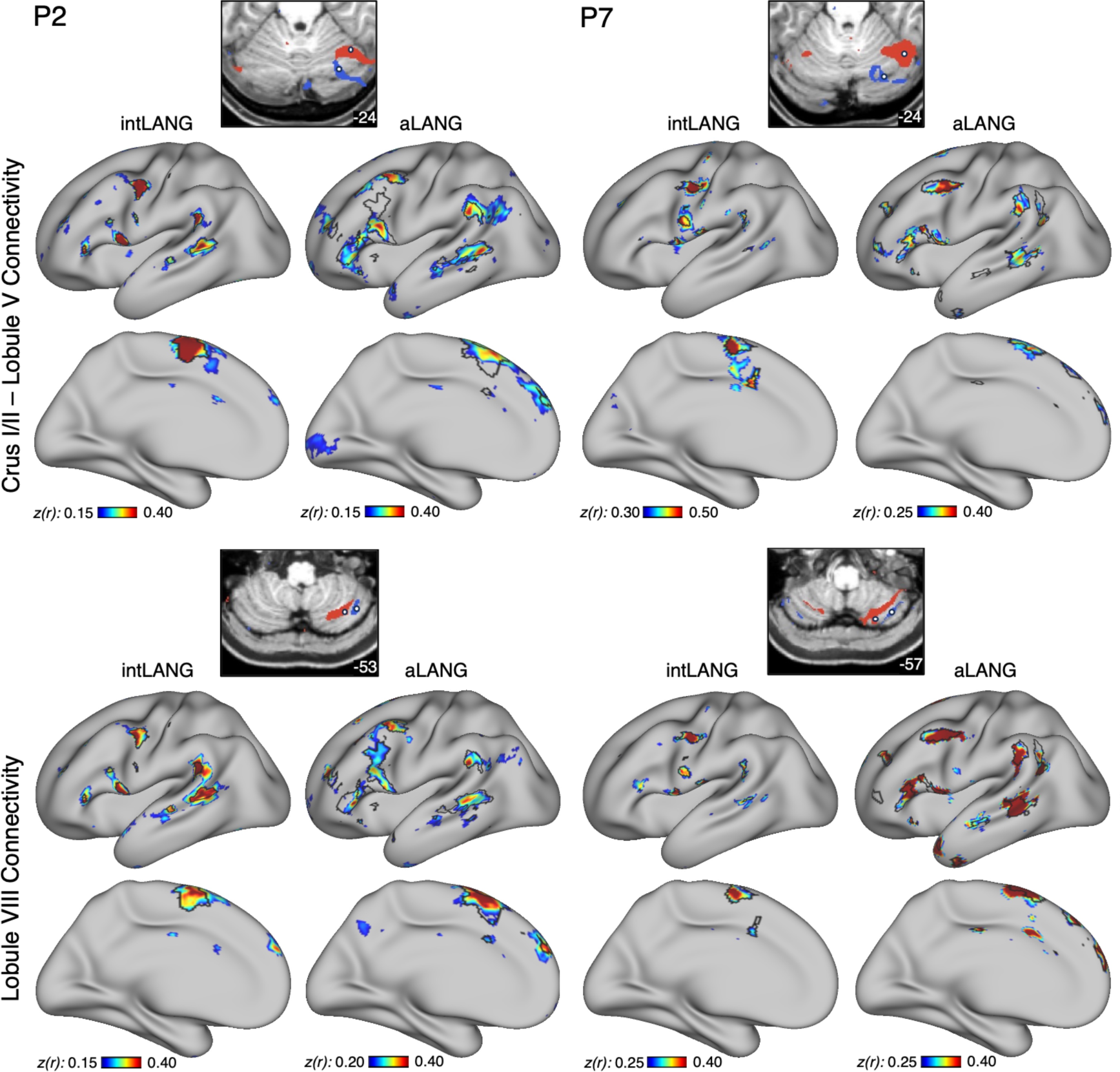
The intermediate language network (intLANG) and association language network (aLANG) are linked to distinct cerebellar regions. Data are shown for two representative participants in the left and right columns, P2 and P7, with the two cerebellar zones displayed in separate rows: the Crus I/II – Lobule V extended zone (**Top rows**) and the Lobule VIII extended zone (**Bottom rows**). Within each zone, a winner-take-all parcellation of the cerebellum, computed from each participant’s Dataset 1 by assigning every cerebellar voxel to the most strongly correlated cerebral network, is displayed on representative cerebellar slices (top of each block), with the intLANG (red) and aLANG (blue) representations occupying spatially segregated regions in each individual. Seed regions were placed within the intLANG and aLANG cerebellar representations, and their functional connectivity across the cerebral cortex was estimated from the independent Dataset 2. The seed-region-based correlation maps are shown on the inflated lateral and medial cortical surfaces for the cerebellar intLANG seed region and the aLANG seed region, with the independently defined cerebral network boundaries overlaid on the surface (black outlines). The intLANG cerebellar seed region recapitulated the distributed cortical intLANG network, anchored at dPCSA, vPCSA, and Spt, whereas the adjacent aLANG cerebellar seed region recapitulated the distinct cortical aLANG network. White circles mark the cerebellar seed region locations. Coordinates in each panel indicate the section level in the MNI152 atlas. Correlation maps are plotted as *z(r)* using the JET256 color scale.

A further detail of the topography is notable. The anterior cerebellar region associated with the intLANG network was positioned between the location of the anterior lobe body map and the major cognitive zones that fall posteriorly within Crus I/II (Saadon-Grosman et al. 2022; 2024). Thus, much like the nested organization in the cerebral cortex, the cerebellar components of intLANG and aLANG display a hierarchical relationship, with intLANG consistently positioned near to known orofacial somatomotor representations.

### intLANG is preferentially recruited by tasks demanding phonological processing

Having established anatomical dissociation between intLANG and aLANG, we next explored their functional response properties, focusing on tasks demanding phonological processing (Hickok 2025). Two available task contrasts probed phonology: a Rhyme > Face working memory contrast from the N-Back task and a Nonword > Fixation contrast from the Sentence Processing task. The first task required active rhyme judgments under working memory load, and the second involved passive reading of pronounceable nonwords devoid of syntactic structure and semantic content.

intLANG was robustly activated for both the Rhyme > Face working memory contrast (Fig. 5A) and the Nonword > Fixation contrast from the Sentence Processing task (Fig. 5B), whereas aLANG showed minimal response. At the regional level (Fig. 5, right columns), this pattern was carried by all three intLANG regions and all five aLANG regions, suggesting that the regions falling within the same network responded similarly. Notably, adjacent regions in the posterior cortex responded differentially: Spt showed an increased response relative to pSTS. Post-hoc statistical analyses confirmed the separation between networks: Rhyme > Face (*t*(4) = 3.72, *p* < 0.05, Cohen’s *d* = 1.66) and Nonword > Fixation (*t*(4) = 3.93, *p* < 0.05, Cohen’s *d* = 1.76). These observations were all replicated in larger participant cohorts using data from Studies 2 and 3.

**Figure 5.**
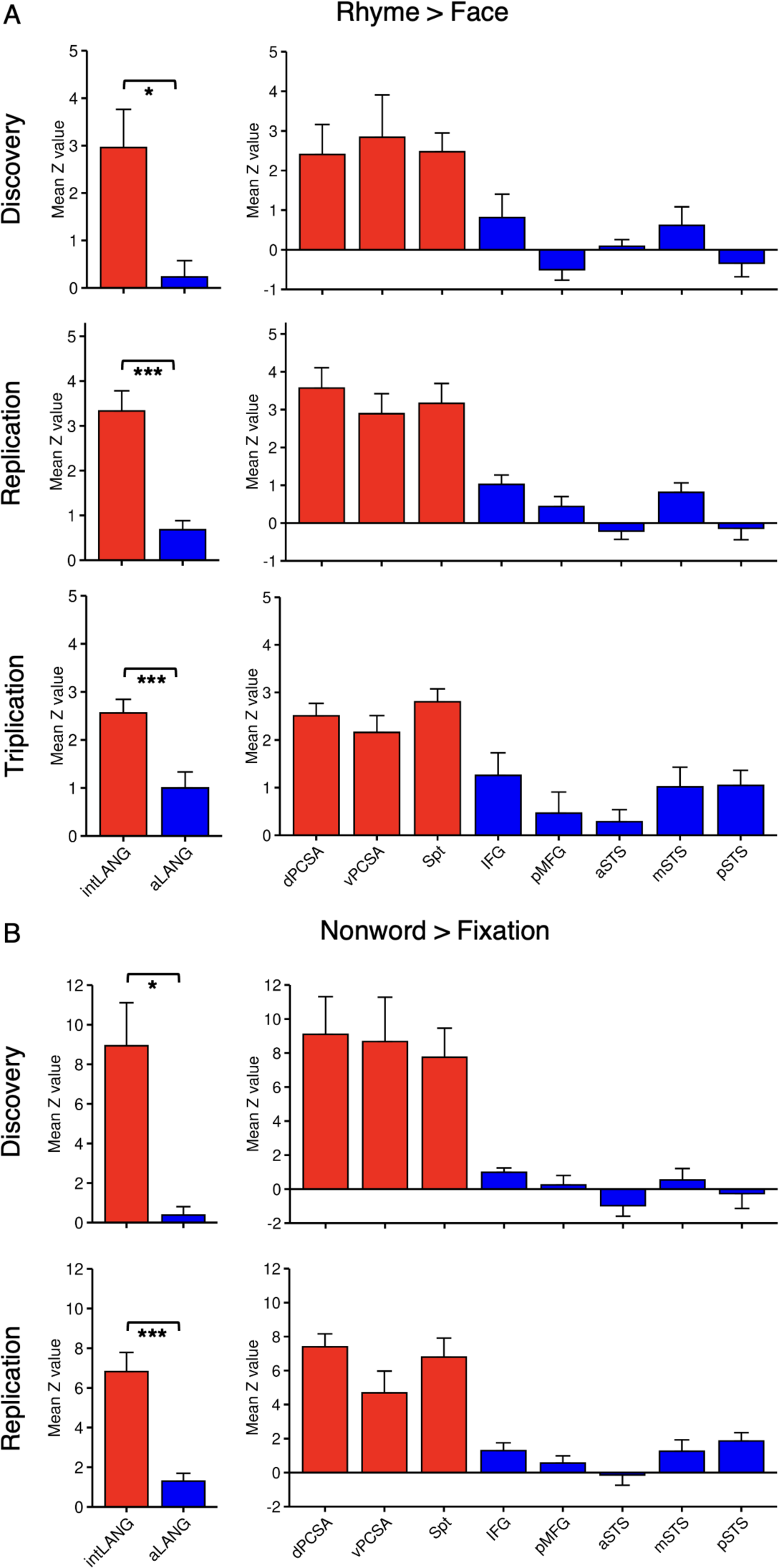
The intermediate language network (intLANG) is preferentially recruited by phonological processing demands. Bar plots show mean functional task response and standard error for two phonological task contrasts within intLANG (red) and the association language network (aLANG, blue), at the network level (**Left bar plots**) and at the individual region level (**Right bar plots**). (**A**) The Rhyme > Face contrast from the N-Back working memory task is plotted for each cohort separately (Studies 1-3, designated Discovery, Replication, and Triplication). intLANG consistently showed robust recruitment while aLANG showed a lesser response. The pattern was found for each of the individual regions within the separate networks: dorsal precentral speech area (dPCSA), ventral precentral speech area (vPCSA), and the Sylvian parietal-temporal area (Spt), each showed a strong positive response, while the five aLANG regions – inferior frontal gyrus (IFG), posterior middle frontal gyrus (pMFG), anterior superior temporal sulcus (aSTS), middle superior temporal sulcus (mSTS), and posterior superior temporal sulcus (pSTS) – showed a smaller response. (**B**) The Nonword > Fixation contrast from the Sentence Processing task is plotted for each cohort separately (Studies 1 and 3, designated Discovery and Replication). The same pattern of preferential intLANG recruitment was observed: intLANG regions (dPCSA, vPCSA, Spt) showed strong positive responses while aLANG regions showed lesser responses. Across both contrasts and all cohorts, intLANG was preferentially recruited by phonological processing. \**p* < 0.05; \*\**p* < 0.01; \*\*\**p* < 0.001. Post-hoc statistical tests of every region individually are reported in Table 1.

**Table 1.** Paired t-tests comparing individual regions of the intLANG and aLANG networks.

| Rhyme > Face<br>intLANG | aLANG |  |  |  |  |
| --- | --- | --- | --- | --- | --- |
|  | *IFG* | *pMFG* | *aSTS* | *mSTS* | *pSTS* |
| *dPCSA* | *** | *** | *** | *** | *** |
| *vPCSA* | *** | *** | *** | *** | *** |
| *Spt* | *** | *** | *** | *** | *** |

| Nonword > Fixation<br>intLANG | aLANG |  |  |  |  |
| --- | --- | --- | --- | --- | --- |
|  | *IFG* | *pMFG* | *aSTS* | *mSTS* | *pSTS* |
| *dPCSA* | *** | *** | *** | *** | *** |
| *vPCSA* | ** | *** | *** | ** | ** |
| *Spt* | *** | *** | *** | *** | *** |

Note: Significance of within-subject paired *t*-tests comparing each intLANG region (rows) against each aLANG region (columns) are shown, pooled across all participants for both the Rhyme > Face contrast (N=22) and the Nonword > Fixation contrast (N=13). All regions were defined *a priori* without examination of the task effects. dPCSA, dorsal precentral speech area; vPCSA, ventral precentral speech area; Spt, Sylvian parietal-temporal area; IFG, inferior frontal gyrus; pMFG, posterior middle frontal gyrus; aSTS, anterior superior temporal sulcus; mSTS, middle superior temporal sulcus; pSTS, posterior superior temporal sulcus. \* $p < 0.05$ ; \*\* $p < 0.01$ ; \*\*\* $p < 0.001$ .

### The anatomical and functional dissociations between intLANG and aLANG replicate

All observations were replicated in independent data. Supplemental Materials show the intLANG and aLANG networks for each participant from Studies 2 and 3. While the spatial locations of the regions within each network varied idiosyncratically, all participants possessed adjacent distributed networks with aLANG surrounding intLANG, paralleling the initial Study 1 exploratory sample.

Region-to-region correlations revealed segregation of intLANG from aLANG, again forming a block-modular pattern (Fig. 3). Statistical tests supported the dissociation: Study 2 within-intLANG *r* = 0.70 (*z(r)* = 0.86), within-aLANG *r* = 0.71 (*z*(r) = 0.89), and between-network *r* = 0.33 (*z(r)* = 0.35) (within-intLANG > between, *t*(8) = 13.42, *p* < 0.001, Cohen’s *d* = 4.47, within-aLANG > between, *t*(8) = 8.15, *p* < 0.001, Cohen’s *d* = 2.72); Study 3 within-intLANG *r* = 0.66 (*z(r)* = 0.79), within-aLANG *r* = 0.74 (*z(r)* = 0.95), and between-network *r* = 0.29 (*z(r)* = 0.30) (within-intLANG > between, *t*(7) = 7.19, *p* < 0.001, Cohen’s *d* = 2.54; within-aLANG > between, *t*(7) = 9.71, *p* < 0.001, Cohen’s *d* = 3.43). Distinct cerebellar representations of intLANG and aLANG were present for each participant, and the separate cerebral networks could be recapitulated by placing adjacent seed regions in the distinct cerebellar zones. Fig. 4 shows one example participant from Study 3 (P7). All participants are displayed in Supplemental Materials. The left-lateralization of intLANG also replicated. In Study 2, intLANG occupied 1.43% of left-hemisphere cortical vertices versus 0.23% of right-hemisphere vertices; the lateralization index averaged 0.75 (0.40 – 1.00). In Study 3, intLANG occupied 1.62% of left-hemisphere cortical vertices versus 0.43% of right-hemisphere vertices; the lateralization index averaged 0.63 (0.30 – 1.00).

Most striking was the reproducibility of the functional dissociations between networks. intLANG displayed a greater response than aLANG to both tasks that demanded phonological processing (Fig. 5). Planned comparisons revealed that the intLANG response exceeded the aLANG response for all three replication datasets (the Rhyme > Face contrast was available for both Studies 2 and 3 and the Nonword > Fixation contrast was available only for Study 3). For the Rhyme > Face contrast, the intLANG response was higher than the aLANG response in Study 2 (*t*(8) = 5.57, *p* < 0.001, Cohen’s *d* = 1.86) and Study 3 (*t*(7) = 5.88, *p* < 0.001, Cohen’s *d* = 2.08). For the Nonword > Fixation contrast, the intLANG response also exceeded the aLANG response in Study 3 (*t*(7) = 7.88, *p* < 0.001, Cohen’s *d* = 2.79). Moreover, across all individual regions there was not a single exception to the preferential pattern of response. That is, each of the three individual regions of intLANG (dPCSA, vPCSA, and Spt) showed a quantitatively greater response than each of the five individual regions of aLANG (IFG, pMFG, aSTS, mSTS, and pSTS).

As a post-hoc analysis, to illustrate the regional consistency of the functional dissociation, data from all participants across the three studies were pooled (N=22 for the Rhyme > Face contrast and N=13 for the Nonword > Fixation contrast) and the comprehensive between-network region-to-region comparisons were tested. Note that these are not independent tests but rather convergent statistical tests to explore whether the omnibus (whole network) effect is also present between each of the included regions. Results are shown in Table 1. In every case, the responses in the individual intLANG regions, independently for both tasks, were significantly greater than the responses in any of the aLANG regions. The differences included closely adjacent anterior and posterior regions, illustrating that the dissociation is predicted by the network affiliations of the regions, not their spatial proximity to one another. For example, the posterior intLANG region, Spt, showed a greater response than the nearby aLANG region, pSTS. The same held for the adjacent pair of anterior regions, with vPCSA’s response exceeding that of IFG.

### intLANG and aLANG functionally dissociate phonological versus higher-order syntactic and semantic processing

The analyses to this point provide convergent evidence that intLANG and aLANG are distinct networks that respond differentially during phonological processing tasks. As a final analysis, we returned to the question of how aLANG, which has been extensively studied using functional localizers that require processing of meaningful sentences, differs from intLANG. We examined two opposing task contrasts available in the Sentence Processing task to explicitly test for a functional double dissociation between the two networks: the Nonword > Fixation contrast that engages phonology, and the Sentence > Nonword contrast that preferentially isolates higher-order syntax and semantics while roughly controlling for phonology. All available data were used (N = 13 with data from Studies 1 and 3).

Results are shown in Fig. 6. Network-level responses were submitted to a 2 (network: intLANG, aLANG) × 2 (contrast: Nonword > Fixation, Sentence > Nonword) repeated-measures ANOVA. The interaction was robust and significant (*F*(1,12) = 25.57, *p* < 0.001), confirming that the two networks responded differentially to the two contrasts. Post-hoc paired *t*-tests revealed that the dissociation was carried by a full crossover interaction: the Nonword > Fixation contrast recruited intLANG over aLANG (*t*(12) = 6.78, *p* < 0.001, Cohen’s *d* = 1.88), whereas the Sentence > Nonword contrast recruited aLANG over intLANG (*t*(12) = 3.19, *p* < 0.01, Cohen’s *d* = 0.88).

**Figure 6.**
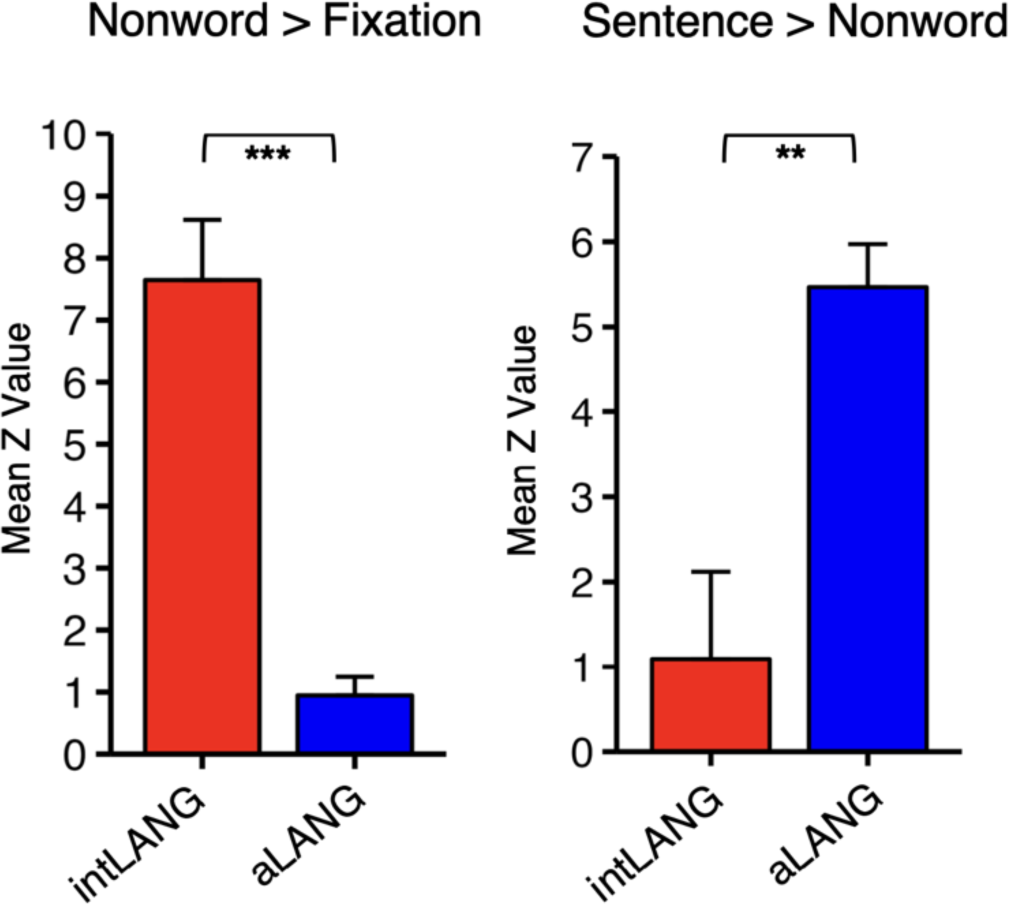
Functional double dissociation between the intermediate language network (intLANG) and the association language network (aLANG). Bar plots show the mean functional task response and standard error within intLANG (red) and aLANG (blue). intLANG showed robust recruitment for the Nonword > Fixation task contrast, while aLANG showed minimal response, consistent with intLANG’s preferential engagement during low-level phonological processing. By contrast, aLANG was strongly recruited by the Sentence > Nonword contrast, while intLANG showed little response, consistent with aLANG’s role in higher-order semantic processing. The interaction was robust (*F*(1,12) = 25.57, *p* < 0.001), confirming that the two networks responded differentially to the two contrasts. Within each contrast, paired *t*-tests between networks supported the observed differences and indicate that the effect is carried by a full crossover interaction. \**p* < 0.05; \*\**p* < 0.01; \*\*\**p* < 0.001.

Thus, two nested networks, intLANG and aLANG, that occupy closely juxtaposed and sometimes interdigitated regions across multiple cortical zones, are preferentially recruited by distinct linguistic dimensions: intLANG by phonology and aLANG by higher-order syntax and semantics.

## Discussion

Human language is supported by at least two nested perisylvian networks that radiate outward from primary motor and auditory cortex. Using within-individual precision functional mapping across three independent studies, we identified an intermediate language network, intLANG, anchored on the precentral speech areas (dPCSA, vPCSA) and Spt, nested within a surrounding association language network, aLANG, that extends into prefrontal and temporal association cortex. The component regions of the two networks are closely juxtaposed across both anterior and posterior cortical language zones, with idiosyncratic spatial variability between individuals. Three lines of evidence establish the two networks as separate.

First, intrinsic connectivity among the component regions displays modular features, exhibiting strong connectivity within each network and weaker connectivity between networks (Fig. 3; see also Braga et al. 2020). Second, the segregation extends beyond the cerebral cortex. Adjacent cerebellar regions recover the distributed cerebral networks in their entirety suggesting that the two networks are components of distinct brain-wide circuits (Fig. 4). Third, the two networks are functionally dissociable: intLANG is preferentially recruited by phonological processing tasks and aLANG by meaning-based sentence processing (Figs. 5 and 6). The phonological dissociation replicated in each study and across all the component distributed regions of each network, including dissociations between adjacent regions in anterior and posterior language zones (Table 1).

These findings argue against the idea that a single core integrative network supports language functions. Rather, language is supported by multiple distributed networks paralleling the broad anatomical motif found in other brain systems, with functionally specialized higher-order networks radiating outward from lower-order networks (Giarrocco and Averbeck 2023; Du et al. 2024; Buckner 2026). We discuss the implications of these findings for language function as well as for understanding the broad anatomical organization and evolution of higher-order cognitive domains.

### The intermediate language network

The anatomical anchors of intLANG, including dPCSA, vPCSA, and Spt, align with regions implicated in the dorsal stream of the dual-stream model of speech processing (Hickok and Poeppel 2007; Hickok 2025). In this framework, a left-lateralized dorsal pathway supports sensorimotor integration for speech, mapping acoustic representations onto articulatory motor representations and providing the feedback signals required for fluent production (Hickok and Poeppel 2007; Hickok 2025; see also Scott and Johnsrude 2003; Rauschecker and Scott 2009; Binder 2015). Among the dorsal-stream regions, area Spt, lying within the posterior Sylvian fissure, has been characterized as a sensorimotor interface for the vocal tract, exhibiting both auditory and motor response properties during covert rehearsal and overt speech production (Warren et al. 2005; Hickok et al. 2009). The present results, along with prior precedents (Glasser et al. 2016; Braga et al. 2020), suggest the component regions form an integrated brain-wide network that is partially segregated from adjacent networks.

The juxtaposition of the network regions, including their interdigitation, may contribute to why intLANG has been difficult to resolve as a distinct distributed network in past functional neuroimaging studies. For example, a recent study examining the response properties of the monolithic language network, identified by a single contrast of meaningful sentences with nonwords, found the network to be sensitive to phonological processing demands, leading the authors to conclude that “the boundaries between different levels of linguistic structure – from phonemes to morphemes to words to constructions and syntactic rules – are not sharp” (Regev et al. 2024). One interpretation of these findings is that the levels of linguistic structure are genuinely not segregated; another, which the present results favor, is that distinct networks supporting different levels of linguistic analysis were merged because their juxtaposed regions were not separated. The present analyses relied on intentional contrasts between functional connectivity patterns to distinguish closely adjacent regions that are otherwise difficult to disentangle (see also Braga et al. 2020). While remaining agnostic about the specific computational content that underlies the response properties of the intLANG network, or fully defining the degree of aLANG involvement across linguistic processing demands, the collective observations provide strong evidence that intLANG is anatomically distinct from aLANG, and differentially specialized for processing phonology in support of language function.

### The higher-order association language network nests around the intermediate language network

The extensively documented language network, which responds during the processing of meaningful sentences, is one of multiple networks that support language function. We label our estimate of this network as the association language network (aLANG), to differentiate it from intLANG while preserving its connection to the network’s characterization by Fedorenko and colleagues (Fedorenko et al. 2024; see also Vandenberghe et al. 2002; Humphries et al. 2006). The nested spatial arrangement between intLANG and aLANG suggests a hierarchy that radiates outward from early sensory and motor cortex. In retrospect, the general features of the nested pattern can be seen in group-averaged maps. For example, in a meta-analysis of 13,500 studies, the term “phonemic” was associated with a broad left-lateralized pattern along the central sulcus extending into auditory regions, while the term ‘semantic’ was linked to a broad adjacent pattern that surrounded it (Voets et al. 2025). The spatial details of the separate network regions are not visible in the meta-analysis, but the broad nesting pattern is observed (Gong et al. 2023). Similarly, in the neurosurgical stimulation meta-analysis of Lu et al. (2021), the frontal region associated with anomia (the inability to name objects) fell anterior to the vPCSA region, within the prefrontal association region linked to aLANG.

Notably, nested arrangements among brain networks are a recurring motif across the cerebral cortex. Outside of the language domain, Giarrocco and Averbeck (2023) proposed a nesting of networks based on anatomical analysis of multiple parietal-frontal pathways in the macaque monkey (see also Averbeck et al. 2009; Xu et al. 2022). In their account, the lowest-level parietal-frontal network begins with the S1 to M1 connections across the central sulcus; this is surrounded by a more distributed network connecting premotor frontal regions with parietal regions just posterior to S1, which in turn is surrounded by an even more distributed parietal-frontal network that extends into canonical prefrontal and parietal association cortex. Margulies and colleagues, drawing on human functional neuroanatomy, noted that gradients of networks radiate outward from early sensory and motor cortex, progressing to distributed association networks that support higher-order cognition at the gradient’s apex (Margulies et al. 2016; Huntenburg et al. 2018). Our laboratory has previously noted that nested networks are a prominent feature of both human and monkey cortical organization (Buckner and Margulies 2019; Buckner and DiNicola 2019; Du and Buckner 2021; Buckner 2026).

The present findings indicate that language networks are a further instance of a hierarchical gradient of networks that begins with early auditory cortex and the orofacial motor representations along the precentral gyrus and radiates outward to the canonical apex zones of association cortex. That specialized language regions might have evolved near precentral motor speech regions and auditory regions important for perception has long been appreciated (Geschwind 1970; Petrides et al. 2005; Krubitzer 2007). What the present results add is that the cortical association zones specialized for higher-order aspects of syntax and semantics constitute a tertiary association network that surrounds the lower-order intLANG network, which in turn surrounds early auditory and motor cortex. We further hypothesize that these two nested networks develop sequentially over the early postnatal years (see Krienen and Buckner 2020; Buckner 2026 for discussion). If language follows this motif, an outward progression – from early sensory and motor cortex to the association apex – may govern both how such faculties arose in evolution and how they assemble in the developing brain (see also Braga et al. 2020; Hickok 2025).

### The multiple networks supporting language are components of segregated brain-wide networks

A canonical circuit connects cerebral networks with subcortical structures including the cerebellum and striatum (Bostan and Strick 2018). The association language network follows this pattern with prominent right-lateralized (contralateral) representations in the cerebellum (Stoodley and Schmahmann 2009; Guell et al. 2018; Xue et al. 2021; Saadon-Grosman et al. 2024; Casto et al. 2026). In the present data, the cerebellum contained separate representations of both the intLANG and aLANG networks, with topographic features that reinforce the hypothesis that the two networks are segregated and form a hierarchy.

In almost every participant, distinct regions associated with both networks could be identified, with representations near to the anterior lobe of the cerebellum and a second pair posteriorly. More revealing are the exact positions and topographic relationships between the two networks within each of the anterior and posterior cerebellar representations. Near the primary fissure that divides the anterior and posterior cerebellar lobes, the intLANG representation lies anteriorly, near the primary body map (e.g., Saadon-Grosman et al. 2022), while aLANG’s representation lies posteriorly, within the higher-order cognitive zones of the cerebellum (Saadon-Grosman et al. 2024). In the second pair of network representations, posterior to the Crus I / II association cluster, the spatial ordering again situates the second intLANG representation near to the second cerebellar body map. This double representation is a consistent feature of cerebellar organization (Buckner 2013; Guell et al. 2018). That the cerebellar aLANG representations lie near to the association megaclusters throughout the cerebellum, while the intLANG representations are consistently situated nearer to the somatomotor body maps, further supports a hierarchical ordering that places intLANG at a lower level and aLANG at the apex. Providing additional evidence that the two networks are segregated, the full distributed extent of each cerebral network could be recapitulated by functional connectivity from small seed regions placed in the adjacent cerebellar regions (Fig. 4).

### Limitations and future directions

The network estimates of intLANG and aLANG were identified using intrinsic functional connectivity, an indirect measure of brain organization (Biswal et al. 1995; Greicius et al. 2003; Fox and Raichle 2007; Van Dijk et al. 2010; Buckner et al. 2013; Power et al. 2014). The findings revealed that the network boundaries predicted task response patterns, supporting the validity of the estimates, but exceptions and boundary mismatches were observed (as they are in all such studies). Convergent efforts that seek to dissociate the network regions using direct electrophysiological recording and electrical stimulation will be an important future direction. Of relevance, in a multicenter neurosurgical study, four spatially distinct anatomical clusters were found that induced speech arrest during direct cortical stimulation (Lu et al. 2021). The anatomical locations of the four clusters correspond roughly with the anatomical regions that comprise the intLANG network, with clusters near to dPCSA, vPCSA, Spt, and also SMA (see also Hickok et al. 2023). Within-individual precision network mapping conducted prior to neurosurgery could allow network estimates to be used directly to localize electrophysiological recording contacts (Sun et al. 2025b) or stimulation sites (Xu et al. 2026).

A second future direction concerns automation of the methods. In the present work, intLANG identification relied on model-free seed-region-based functional connectivity, which requires no assumptions to interpret the network estimates. In our experience, this is how novel network discoveries have generally been first made (e.g., Vincent et al. 2006; Braga and Buckner 2017; see also Gordon et al. 2023 for a recent example). To be applied routinely, however, particularly in translational settings, more automated approaches are desirable. Our attempts to recover intLANG using model-based approaches have so far been unsuccessful. Two features contribute. First, intLANG occupies a small cortical territory anchored to compact regions and, compared with larger association networks, is at greater risk of being absorbed into its neighbors when group priors are applied. Second, intLANG sits in juxtaposition with multiple distinct networks whose boundaries vary idiosyncratically, which can cause intLANG to be misassigned or split across them. Future work developing model-based methods capable of recovering small, compact networks, or hybrid approaches combining seed-region-based initialization with model-based refinement, will be valuable for scaling intLANG identification to larger samples.

How the multiple nested networks interact is another important question for future research. The present analyses focused on dissociations and so emphasize separations. We expect that our coarse methods underemphasize regions of overlap, as well as subdivisions within the language zones that fall below the resolution of current functional neuroimaging methods. Further progress may require scanning at higher resolution, at 7T or above, or combining functional neuroimaging with direct neurophysiological recordings. There are, nonetheless, substantive follow-up explorations that can be undertaken with current technologies. For example, while we emphasized network differences, some regions overlapped between the two networks. Some of this overlap, and perhaps most of it, may reflect mundane spatial blurring given the limited resolution of the methods. But some may genuinely reflect regions of interaction between the two networks – candidate hubs of convergence (see Ladwig et al. 2026 for recent discussion). One such candidate was noted in IFG at or near classically defined Broca’s area. Comprehensively characterizing these candidate regions of convergence will be an interesting forward direction.

A final limitation is that the Sentence Processing task used in the present study was originally designed as a localizer for the association language network (Fedorenko et al. 2010) and was not optimized to target intLANG specifically. We repurposed the Nonword > Fixation contrast as a phonological probe on the rationale that pronounceable nonwords engage grapheme-to-phoneme conversion without lexical-semantic content. The contrast revealed strong intLANG recruitment and converged with an independent Rhyme > Face contrast from the N-Back working memory task. Characterizing intLANG’s response properties more precisely will require task paradigms designed specifically to manipulate phonological processing demands and contrast them with alternatives, such as those developed by Hickok and colleagues (discussed extensively in Hickok 2025), Scott and Perrachione (2019), or applied by Regev et al. (2024). A key requirement will be to dissociate phonological processing itself from the articulatory rehearsal, working memory, and segmentation demands that metalinguistic tasks typically require, since these associated operations, rather than phonological representation per se, may account for much of the recruitment observed in such paradigms. Overall, the present results provide strong evidence for anatomical separation between the networks but do not survey or manipulate linguistic processes sufficiently to specify the detailed processing contributions of each.

## Conclusions

Language is supported by partially segregated, functionally differentiated networks organized in a nested arrangement that radiates outward from orofacial motor regions and the early auditory regions. Here we characterized and dissociated two of these networks: intLANG, an intermediate network biased toward phonology, and a surrounding association network, aLANG, that subserves higher-order syntax and semantics. This nested organization echoes the arrangement of other brain systems, suggesting a shared hierarchical motif that may have evolved to support specialized higher-order cognitive functions across multiple gradients.

## Methods

### Participants and MRI data acquisition

Details of the MRI methods for Studies 1 and 3 have previously been reported in Du et al. (2024) and for Study 2 in Sun et al. (2025a). Relevant portions are repeated here.

### Participants

Across studies, 22 English-speaking volunteers participated for payment. All were native English speakers except two, who were fluent English speakers from the age of 6. Participants provided written informed consent under protocols approved by the Institutional Review Boards of Harvard University (Studies 1 and 3) and Massachusetts General Hospital (Study 2). Study 1 participants (P1–P5, N = 5; 3 women) and Study 3 participants (P6–P15, N = 8; 6 women) were a subset of those reported in Du et al. (2024). Two of the original 10 participants (P10 and P11) were excluded due to insufficient task data; the remaining participant IDs were preserved to maintain consistency with prior reports. Study 2 participants (S1–S9, N = 9; 3 women) were an entirely separate cohort from Sun et al. (2025a). While Sun et al. (2025a) included a patient cohort, only control participants were analyzed here. All participants were right-handed.

### Scanner and setup

Scanning was performed at the Harvard Center for Brain Science using a 3T Siemens MAGNETOM Prisma^fit^ MRI scanner and a 32-channel phased-array head-neck coil (Siemens Healthineers AG, Erlangen, Germany). Foam and inflatable padding were used to minimize head motion. Participants viewed a rear-projected display positioned to optimize comfortable viewing. Eyes were video recorded using an Eyelink 1000 Core Plus with Long-Range Mount (SR Research, Ottawa, Ontario, Canada). MRI data quality was monitored during the scan using Framewise Integrated Real-time MRI Monitoring (FIRMM; Dosenbach et al. 2017) for Studies 1 and 3 and using ScanBuddy for Study 2 (Asay et al. 2025).

### Functional data acquisition

Across all studies, blood oxygenation level-dependent (BOLD) data were acquired using identical sequence parameters: voxel size = 2.4 mm, TR = 1,000 ms, TE = 33.0 ms, flip angle = 64°, matrix 92 × 92 × 65 (FOV = 221 × 221 mm), 65 slices covering the full cerebrum and cerebellum. Each resting-state fixation run lasted 7 min 2 sec (422 frames; first 12 frames removed for T1 equilibration). Dual-gradient-echo B0 fieldmaps were acquired (TE = 4.45, 6.91 ms; slices matched to the BOLD sequence). For Studies 1 and 3, each participant was scanned across 8–11 sessions, most often over 6 to 10 weeks (with a few longer gaps up to one year between first and last sessions), yielding 16–24 resting-state fixation runs and 48–70 task-based runs per individual. For Study 2, each participant completed a baseline session containing 8 resting-state fixation runs (with a 10–15 min break midway) and 10 subsequent fMRI sessions distributed over 5 days (2 sessions per day, separated by at least 4 days). Each subsequent session contained one resting-state fixation run (always second), two N-Back task runs, and two Oddball task runs, with the order of the two task types counterbalanced across sessions, yielding 16-18 resting-state fixation runs and 36-40 task-based runs per individual. Three field maps were acquired during the baseline session and two during each subsequent session.

### Structural data acquisition

For Studies 1 and 3, high-resolution T1w and T2w structural images were acquired using Human Connectome Project (HCP) sequences (Harms et al. 2018). T1w magnetization-prepared rapid gradient echo (MPRAGE) parameters: voxel size = 0.8 mm, TR = 2,500 ms, TE = 1.81/3.60/5.39/7.18 ms, TI = 1,000 ms, flip angle = 8°, matrix 300 × 320 × 208, 208 slices, in-plane generalized auto-calibrating partial parallel acquisition (GRAPPA) acceleration = 2. T2w sampling perfection with application-optimized contrasts using different flip angle evolution sequence (SPACE) parameters: voxel size = 0.8 mm, TR = 3,200 ms, TE = 564 ms, matrix 300 × 320 × 208, 208 slices, in-plane GRAPPA acceleration = 2. For Study 2, high-resolution T1w and T2w images were acquired using FreeSurfer sequences during the baseline session. T1w MPRAGE: voxel size = 1.0 mm, TR = 2,530 ms, TE = 1.69/3.55/5.41/7.27 ms, TI = 1,100 ms, flip angle = 7°, matrix 256 × 256 × 192, 192 slices, in-plane GRAPPA acceleration = 2. T2w SPACE: voxel size = 1.0 mm, TR = 3,200 ms, TE = 564 ms, matrix 256 × 256 × 192, 192 slices, in-plane GRAPPA acceleration = 2. For all studies, backup rapid T1w structural images were obtained using a multi-echo MPRAGE sequence (van der Kouwe et al. 2008). For Studies 1 and 3: voxel size = 1.2 mm, TR = 2,200 ms, TE = 1.57/3.39/5.21/7.03 ms, TI = 1,100 ms, flip angle = 7°, matrix 192 × 192 × 144, in-plane GRAPPA acceleration = 4; for Study 2: voxel size = 1.2 mm, TR = 2,200 ms, TE = 1.57/3.35/5.13/6.91 ms, TI = 1,100 ms, flip angle = 7°, matrix 210 × 210 × 192, in-plane GRAPPA acceleration = 4.

### MRI data processing

Data were processed using the openly available iProc pipeline (Braga et al. 2019). iProc was designed to maximize cross-session alignment within individuals while minimizing interpolations to preserve the detailed functional anatomy of each participant. For Studies 1 and 3, processed resting-state and task-based data were taken directly from Du et al. (2024) and Du et al. (2025). For Study 2, data from Sun et al. (2025a) were processed using the same iProc pipeline. Relevant details are summarized below; minor procedural differences arose between the cohorts because Sun et al. (2025a) included a patient cohort in the larger sample that necessitated more stringent motion correction (e.g., including timepoint censoring for motion which was not performed for the Du et al. 2024 participants due to uniform low motion).

Data were interpolated to a 1-mm isotropic T1w native-space atlas, with all processing steps composed into a single interpolation, and then projected using FreeSurfer v6.0.0 to the fsaverage6 cortical surface (40,962 vertices per hemisphere; Fischl et al. 1999). Four transformation matrices were calculated: (1) a motion correction matrix for each volume to the run’s middle volume (linear registration, 6 degrees of freedom [DOF]; MCFLIRT, FSL; Jenkinson et al. 2002); (2) a matrix for field-map-unwarping the run’s middle volume, correcting for field inhomogeneities caused by susceptibility gradients (FUGUE, FSL; Jenkinson et al. 2012); (3) a matrix registering the field-map-unwarped middle BOLD volume to the within-individual mean BOLD template (12 DOF; FLIRT, FSL; Jenkinson and Smith 2001); and (4) a matrix registering the mean BOLD template to the participant’s T1w native-space image, which was resampled to 1.0-mm isotropic resolution (6 DOF; boundary-based registration, FreeSurfer). The individual-specific mean BOLD template was created by averaging all field-map-unwarped middle volumes after registering them to an upsampled 1.2-mm unwarped mid-volume template (an interim target, selected from a low-motion run typically acquired close to a field map).

For resting-state fixation and task-based runs used for functional connectivity analysis, confounding variables including six head motion parameters, whole-brain, ventricular and deep cerebral white matter signals, and their temporal derivatives were calculated from the BOLD data in T1w native space and regressed out using 3dTproject (AFNI; Cox 1996, 2012). The residual BOLD data were then bandpass filtered at 0.01–0.1 Hz using 3dBandpass (AFNI; Cox 1996, 2012). For runs used for standard task contrast analysis, only whole-brain signal was regressed out (see DiNicola et al. 2020). The data were then resampled from T1w native-space atlas to the fsaverage6 cortical surface mesh using trilinear interpolation and surface-smoothed using a 2-mm full-width-at-half-maximum (FWHM) Gaussian kernel.

For Study 2, confound regression used an expanded 36-parameter set (six head motion parameters, three global tissue signals, their temporal derivatives, and the quadratic terms of all 18; Ciric et al. 2017). Volumes with high framewise displacement (>0.4 mm or >3 SD above the baseline-session mean) were additionally flagged and added to the nuisance regression matrix; for one participant with respiratory-related oscillations, the threshold was raised to 0.7 mm (∼3 SD above baseline). All other processing steps were identical to those in Studies 1 and 3.

### Identification of intLANG and its relations to adjacent networks within individuals

intLANG was identified within each individual using model-free seed-region-based intrinsic functional connectivity, following procedures adapted from Braga and Buckner (2017) and Braga et al. (2019). The specific procedure for identifying the two segregated language networks was fully developed in the Study 1 participants and then prospectively applied to the Study 2 and 3 participants without modification. Furthermore, each participant’s extensive data were divided into independent Datasets 1 and 2 (Supplemental Materials). In this manner, all effects could be replicated across independent participant cohorts as well as within each participant’s idiosyncratic anatomy.

### Functional connectivity matrices

Functional connectivity matrices were generated for each participant individually and included independent (replicate) datasets, referred to as Dataset 1 and Dataset 2. Functional connectivity was initially computed on the cortical surface. For each run, pairwise Pearson correlations were calculated between the BOLD time series at every pair of cortical vertices, producing an 81,924 × 81,924 vertex-wise correlation matrix (40,962 vertices per hemisphere). Correlations were Fisher *z*-transformed and averaged across two separate (split-half) datasets within each individual (Dataset 1 and Dataset 2), and inverse-Fisher-transformed back to *r* values to yield stable within-individual mean correlation matrices. The split into within-individual Dataset 1 and Dataset 2 excluded data from the Sentence Processing task, reserved for independent functional characterization. The matrices were assigned to the vertices of an in-house cortical template (Braga and Buckner 2017), enabling interactive selection of single-vertex seeds and real-time visualization of the resulting correlation maps in Connectome Workbench’s wb_view (Marcus et al. 2011). The number of runs included in the datasets for each individual is reported in Supplemental Materials. Note that the extensive, repeat scanning across sessions within individuals allowed each of the replicate datasets (i.e., Dataset 1 and Dataset 2) to each independently contain a very large amount of data. These correlation matrices were used both to explore manual seed regions placed at individual vertices to identify intLANG and for the subsequent extraction of regional-level time series for targeted analyses. The presence of independent datasets allowed networks and regions to be developed in one dataset and then prospectively applied to the replicate, independent datasets.

### Identification of intLANG

intLANG was identified within each individual using seed regions manually placed in Dataset 1. The tongue / mouth motor representation was first localized, using the Tongue > All task contrast where somatomotor task data were available and seed-region-based functional connectivity otherwise (see Tongue motor localizer below). Single-vertex seed regions were then placed along the precentral gyrus surrounding that representation, and the resulting correlation maps were examined for evidence of a distributed network. These initial seeds revealed a left-lateralized network comprising dorsal and ventral precentral regions (dPCSA and vPCSA), a posterior region at the parietal-temporal junction (Spt), and a midline region at or near SMA that was spatially distant from the three lateral regions and had not been anticipated. To establish that these separated regions belong to a single network, single-vertex seed regions were subsequently placed within each component in turn, dPCSA, vPCSA, Spt, and SMA, and the resulting correlation maps compared.

The BOLD time series was extracted from each seed region vertex and correlated with all other cortical vertices, yielding a seed-region-based correlation map. Correlation maps were thresholded on a per-participant basis (reported in Supplemental Materials). To illustrate the reproducibility of the network, the seed region vertex identified in Dataset 1 was applied to Dataset 2 to test the reliability of the network correlation pattern. As will be revealed, at this amount of data per participant, the network patterns identified in Dataset 1 were reproduced with an extremely high level of spatial specificity in every individual.

### Identification of adjacent networks

aLANG, AMN, and the tongue motor region (labeled Tongue) were defined within each individual to characterize how intLANG is situated relative to its neighbors. aLANG and AMN were estimated using the same model-free seed-region-based functional connectivity procedure used for intLANG. For each participant, single-vertex seed regions were placed in each network following the anatomical conventions of Du et al. (2024), and the resulting correlation maps were thresholded on a per-individual basis to recover each distributed network. The Tongue region within the ventral portion of the posterior precentral gyrus was defined from the Tongue > All task-based contrast (Saadon-Grosman et al. 2022). For participants without motor task data, the Tongue region was estimated using the same seed-region-based functional connectivity procedure. The same procedures were applied across Studies 1–3.

### Anchor region definitions

Anchor regions were defined within each individual for both intLANG and aLANG and used for analyses of within-and between-network correlation and task response. In each case, regions were defined in Dataset 1 and the measures of interest extracted from the independent Dataset 2, or from the task data, ensuring estimates were unbiased. The three intLANG anchor regions were dPCSA, vPCSA, and Spt, following the general conventions of Hickok et al. (2003, 2023). The five aLANG anchor regions were IFG, pMFG, aSTS, mSTS, and pSTS (see Braga et al. 2020). Each region was defined by first drawing a generous border on that individual’s inflated surface in Connectome Workbench to delineate the anatomical zone of interest, converting the border to a mask with wb_command, and then intersecting that mask with the network estimate from the same individual’s Dataset 1. The final anchor region definitions therefore comprised only those vertices assigned to that individual’s own network estimate within the drawn zone. All anchor regions for all individuals are illustrated in Supplemental Materials.

### Functional connectivity within and between intLANG and aLANG

To test whether intLANG and aLANG are distinct networks rather than components of a single distributed network, we examined the pattern of intrinsic functional connectivity among the regions of each network. The logic is straightforward: if the two networks are functionally segregated, regions within a network should be more strongly correlated with one another than with regions of the other network.

For each individual, the three intLANG anchor regions (dPCSA, vPCSA, and Spt) and five aLANG anchor regions (IFG, pMFG, aSTS, mSTS, and pSTS) defined above (see *Anchor region definitions*) were used, with regions defined in Dataset 1 and connectivity estimated from the independent Dataset 2, ensuring an unbiased estimate. The mean BOLD time series for each region was extracted from Dataset 2, and pairwise Pearson correlations were computed among the time courses of all eight regions for each run, then Fisher *z*-transformed and averaged across runs within each individual to obtain a stable estimate of functional connectivity. Individual region-to-region matrices were then averaged across individuals within each cohort and converted back to *r* values for visualization.

To visualize the region-to-region correlation patterns, matrices were arranged with intLANG regions in the upper-left 3 x 3 block and aLANG regions in the lower-right 5 x 5 block, so that the two off-diagonal blocks index between-network connectivity. For each individual, we computed the mean within-intLANG, within-aLANG, and between-network correlations, and tested whether within-network connectivity exceeded between-network connectivity using a paired *t*-test across participants. The same procedure was applied independently to each of the three studies to assess reproducibility. As a post-hoc analysis, the data from all studies were combined and the segregation effect was explored for each region individually, testing whether its correlation strength was stronger within its own network as compared to every region in the other network.

### Task paradigms and analysis

#### Task-based fMRI data

Extensive task-based fMRI data were analyzed. Analyses for Studies 1 and 3 were taken from Du et al. (2024), with relevant portions of the methods repeated here. Study 2 was analyzed similarly, with the small differences in methods noted below. Within all participants, the same BOLD sequence was used for both task and resting-state fixation runs, ensuring spatial alignment of the estimated networks from both data types within each individual. Task runs served two roles in the present work. First, with stimulus-evoked task structure regressed out, task runs were pooled with resting-state fixation runs to maximize the amount of data available for network estimation (Du et al. 2025) and for independent split-half analyses within individuals. Second, task contrasts from the N-Back working memory task (Studies 1-3) and the Sentence Processing task (Studies 1 and 3) were used to characterize the response properties of intLANG and aLANG. Sentence Processing runs were excluded from the network estimation pool to ensure an independent double dissociation between intLANG’s recruitment by low-level phonological processing and aLANG’s recruitment by high-level semantic processing.

#### N-Back working memory task

The goal of the present analyses was to test whether intLANG was preferentially recruited by phonological processing. For Studies 1 and 3, each run contained four pairs of 25-s task blocks interleaved with 15-s extended fixation blocks. Each block presented one combination of working memory load (0-Back or 2-Back) and stimulus category (Face, Scene, Letter, or Rhyme). All Study 1 and 3 participants completed eight runs of the task. Each run lasted 4 min 44 s (284 frames; the first 12 frames removed for T1 equilibration). For Study 2, each run lasted 3 min 27 s (207 frames; the first 12 frames removed for T1 equilibration) and contained four 30-s task blocks interleaved with 15-s extended fixation blocks. The working memory load of all blocks in Study 2 was 2-Back and stimulus categories included only Face and Rhyme. All Study 2 participants completed 20 runs and had at least 18 usable task runs.

For the present analyses, which targeted phonology, we focused on the Rhyme versus Face task contrast. Rhyme stimuli were 1-syllable English words; in the Rhyme condition, a correct match was a rhyming word (e.g., “treat” matched “street”), making the Rhyme condition a phonological-matching task. Face stimuli were unfamiliar faces from the HCP (Barch et al. 2013) in Studies 1 and 3 and color images generated with StyleGAN (Karras et al. 2019) in Study 2, with an equal representation of male and female faces and diversity across racial phenotypes. In the Face condition, a correct match was an identical face. The Face condition served as a non-linguistic visual control roughly (but imperfectly) matched for working memory and motor demands but lacking phonological content. Each block began with a cue and contained 10 stimuli (2 s each for Studies 1 and 3 and 2.5 s for Study 2, 0.5 s inter-trial interval for all studies). Participants responded “Match” (right index finger) or “No-Match” (left index finger) on each trial following the cue. Block order was counterbalanced across runs such that no category appeared twice in a row in Studies 1 and 3, and such that an equal number of Face and Rhyme blocks appeared in each block position in Study 2. Full details of lure structure, counterbalancing, and load manipulation for Studies 1 and 3 are provided in Du et al. (2024). For Study 2, all blocks included two target and two lure (repeated nontarget) trials. Targets and lures were equally likely to appear in each viable trial position within and across runs. Target and lure orders were counterbalanced across runs and experimental days. Category orders were set for each experimental day, with equal numbers of Face and Rhyme blocks in each block position, and the same category order maintained between days.

#### N-Back task analysis

Task data were analyzed with the general linear model (GLM) as implemented by FSL’s first-level FEAT (FSL v5.0.4), using a canonical double-gamma hemodynamic response function and its temporal derivative. Each block (including the cue) was modeled as a separate regressor. For each participant, we computed first-level contrast maps for the Rhyme and Face conditions (collapsed across 0-Back and 2-Back for Studies 1 and 3, and 2-Back only for Study 2), as well as the Rhyme > Face contrast, which served as the primary dependent measure. intLANG and aLANG network estimates, defined a priori from intrinsic functional connectivity, were applied to extract the mean *z*-value within each network for each contrast in each individual.

#### Sentence Processing task

The Sentence Processing task was adapted from Fedorenko et al. (2010, 2012). The goal of the present analyses was to dissociate the functional response of aLANG and intLANG. This task contrasts visually presented sentences with length-matched nonword strings. Participants passively read sentences (e.g., “IN THE MORNING THE TAILOR WAS SHOWING DIFFERENT FABRICS TO THE CUSTOMER”) or pronounceable nonword strings (“SMOLE MUFRISONA VEDER SMOP FO BON FE PAME OMOSTREME GURY U FO”). Stimuli were presented one word (or nonword) at a time (0.45 s per word). Each 6-s sentence or nonword string was followed by a 0.5-s cue prompting participants to make a right index finger button press, ensuring sustained attention. No stimulus was repeated across the experiment. Sentence and nonword strings were presented in 18-s blocks of three strings each. Eight task blocks per run were interleaved with 18-s extended fixation blocks placed at the start of each run and after every fourth task block. Each run lasted 5 min 0 s (300 frames; the first 12 frames removed for T1 equilibration).

#### Sentence Processing task analysis

The Sentence Processing task was originally designed as a language network localizer focusing on the network we label in this paper as aLANG (Fedorenko et al. 2010) and not constructed to target intLANG. Nonetheless, the task supports two complementary contrasts that target the two networks separately. The Sentence > Nonword contrast preferentially isolates high-level syntactic and semantic processing. Nonword > Fixation is a broader contrast and includes phonological processing as well as attentional and visual processing. Although this contrast is not selective for phonology, intLANG is anatomically situated in precentral and superior temporal regions, well outside early visual and typical posterior attention control regions, so the response extracted within the intLANG mask can be reasonably assumed to reflect phonological rather than visual contributions. Pronounceable nonwords engage grapheme-to-phoneme conversion in the absence of lexical-semantic content, providing a targeted phonological probe. We therefore used Sentence > Nonword as the contrast to functionally dissociate aLANG from intLANG, and Nonword > Fixation as the contrast to dissociate intLANG from aLANG.

#### Tongue motor localizer

The somatomotor task was extended from Buckner et al. (2011) and previously described in Saadon-Grosman et al. (2022). The full task examined foot, glute, hand, and tongue representations. For the present study, only the tongue condition was used to identify the spatial location of an orofacial motor effector region along the precentral gyrus. Briefly, participants performed alternating tongue movements from right to left (touching the premolar upper teeth) in 10-s movement blocks paced by a slow-flickering black circle (right movement cued by circle onset; left movement cued by circle offset). Each run contained four tongue blocks interleaved with blocks of other movement types and 16-s passive fixation blocks. Each run lasted 7 min 8 s (428 frames; the first 12 frames removed for T1 equilibration). Six runs were collected per participant with fully counterbalanced movement orders. Runs were excluded if participants missed or failed to respond to cues. Full task design parameters are detailed in Du et al. (2024).

#### Tongue motor localizer analysis

The Tongue > All contrast, in which the tongue condition was contrasted against the average of all other movement conditions (following Saadon-Grosman et al. 2022), served as the localizer in each individual. For participants without usable somatomotor task data (P1 and P5), the region was localized using seed-region-based functional connectivity anchored in the inferior portion of the precentral gyrus within each participant’s idiosyncratic anatomy. The resulting correlation map recovered the approximate location of the somatomotor tongue region. All conclusions about spatial relations were made based on the directly mapped subset of participants.

### Cerebellar representations of intLANG and aLANG

We tested for the presence of cerebellar representations of intLANG and aLANG, and further tested whether seed-region-based analyses of nearby locations within the cerebellum could reproduce the full distributed extents of the intLANG and aLANG networks (using procedures adapted from Xue et al. 2021 and Saadon-Grosman et al. 2024). This analysis used the same within-individual replicate datasets (Dataset 1 and Dataset 2) used for the cortical surface analyses, but additionally used within-individual data in volume format.

Cerebellar voxels were analyzed in the volume space (MNI152 atlas). The cerebellum was defined by an oversized (generous) cerebellar mask created by dilating a cerebellar mask generated through FreeSurfer’s “recon-all” using a disc with a radius of six voxels (Saadon-Grosman et al. 2024). The candidate networks for the winner-take-all assignment were taken from a single cortical parcellation constructed for each participant. intLANG and aLANG were defined from the within-individual estimates of the present analysis, and all remaining networks were taken directly from the partcipant’s within-individual cortical network estimates in Du et al. (2024), estimated with a multi-session hierarchical Bayesian model (MS-HBM; Kong et al. 2019). Vertices belonging to the intLANG and aLANG estimates were relabeled accordingly, with all other vertices retained their original MS-HBM label. Each cortical vertex therefore carried exactly one network label. We first calculated functional connectivity between the cerebellar voxels and the cerebral networks. Specifically, for each run of Dataset 1, we calculated Pearson’s correlation between the time course of every cerebellar voxel and the mean time course of each cortical network. The resulting connectivity values were Fisher *z*-transformed and averaged across all runs within each participant. No spatial priors were applied in assigning the cerebellar voxels to cerebral networks, and each voxel could be assigned to any cortical network. The network assigned to each cerebellar voxel was the cortical network with which its time course was most strongly correlated (Saadon-Grosman et al. 2024).

To explore the relation between the cerebellar and cerebral cortices without strong assumptions, we undertook a second analysis using a model-free seed-region-based approach (Saadon-Grosman et al. 2024). For these analyses, single-voxel seed regions were placed within the intLANG and aLANG representations in the winner-take-all parcellation of the cerebellum, and the unconstrained functional connectivity patterns across the cerebral cortex were derived in the independent Dataset 2 for each participant and visualized in relation to the cortical network boundaries. A confirmation of the cerebellar network estimate would be established if a cerebellar seed region reproduced the corresponding network’s pattern in the cerebral cortex. By contrast, if the cortical correlation maps were nonspecific, spanned multiple networks, or fractionated networks, then the assignment would be undermined. Because the parcellation and the seed-region-based analysis were derived from independent split-halves, this analysis served as a cross-validated control check.

### Quantification of hemispheric lateralization

To quantify the hemispheric lateralization of intLANG, we used two complementary metrics (Braga et al. 2020). For each individual, the number of cortical vertices assigned to intLANG in each hemisphere was counted using the same per-individual *z(r)* threshold applied during network identification, with medial wall vertices excluded from all counts. First, we computed the percentage of total cortical vertices assigned to intLANG within each hemisphere, providing a measure of network occupancy per hemisphere. Second, we computed a lateralization index for each individual: LI = (L − R) / (L + R), where L and R denote the number of intLANG vertices in the left and right hemispheres, respectively (Binder et al. 1995; Mahowald and Fedorenko 2016; see also Braga et al. 2020). LI ranges from −1 (fully right-lateralized) to 1 (fully left-lateralized), with 0 indicating no hemispheric asymmetry.

## Software and Code Availability

Functional connectivity was computed as Pearson product-moment correlations in MATLAB (v2019a; MathWorks, Natick, MA). Image preprocessing was carried out using a combination of FreeSurfer v6.0.0, FSL, and AFNI. Cortical surface maps were generated in Connectome Workbench v1.3.2, which was also used for model-free seed region verification. Statistical analyses were conducted in R v4.4.1.

## Data Availability

Individual participant data are available through the NIH repository (https://nda.nih.gov). Task descriptions, contrast descriptions, and analysis code are provided on Harvard Dataverse (https://doi.org/10.7910/DVN/AVB4BW).

## Acknowledgments

Evelina Fedorenko generously provided the stimuli and inspiration to characterize the aLANG network. Discussion with Edward Chang inspired our explorations of the intLANG network. We thank the Harvard Center for Brain Science neuroimaging core and FAS Division of Research Computing for their support. We thank Tim O’Keefe for assistance in optimization of data processing, Abbey Russell for assistance with data analysis, and Ross Mair for MRI physics support. The multi-band EPI sequence was generously provided by the Center for Magnetic Resonance Research (CMRR) at the University of Minnesota.

